# The evolution of CHROMOMETHYLASES and gene body DNA methylation in plants

**DOI:** 10.1101/054924

**Authors:** Adam J. Bewick, Chad E. Niederhuth, Ji Lexiang, Nicholas A. Rohr, Patrick T. Griffin, Jim Leebens-Mack, Robert J. Schmitz

## Abstract

**Background:** The evolution of gene body methylation (gbM), its origins and its functional consequences are poorly understood. By pairing the largest collection of transcriptomes (>1000) and methylomes (77) across Viridiplantae we provide novel insights into the evolution of gbM and its relationship to CHROMOMETHYLASE (CMT) proteins.

**Results:** CMTs are evolutionary conserved DNA methyltransferases in Viridiplantae. Duplication events gave rise to what are now referred to as CMT1, 2 and 3. Independent losses of CMT1, 2 and 3 in eudicots, CMT2 and ZMET in monocots and monocots/commelinids, variation in copy number and non-neutral evolution suggests overlapping or fluid functional evolution of this gene family. DNA methylation within genes is widespread and is found in all major taxonomic groups of Viridiplantae investigated. Genes enriched with methylated CGs (mCG) were also identified in species sister to angiosperms. The proportion of genes and DNA methylation patterns associated with gbM are restricted to angiosperms with a functional CMT3 or ortholog. However, mCG-enriched genes in the gymnosperm *Pinus taeda* shared some similarities with gbM genes in *Amborella trichopoda*. Additionally, gymnosperms and ferns share a CMT homolog closely related to CMT2 and 3. Hence, the dependency of gbM on a CMT most likely extends to all angiosperms and possibly gymnosperms and ferns.

**Conclusions:** The resulting gene family phylogeny of CMT transcripts from the most diverse sampling of plants to date redefines our understanding of CMT evolution and its evolutionary consequences on DNA methylation. Future, functional tests of homologous and paralogous CMTs will uncover novel roles and consequences to the epigenome.

## BACKGROUND

DNA methylation is an important chromatin modification that protects the genome from selfish genetic elements, is important for proper gene expression, and is involved in genome stability. In plants, DNA methylation is found at cytosines (C) in three sequence contexts: CG, CHG, and CHH (H is any nucleotide, but G). A suite of distinct *de novo* and maintenance DNA methyltransferases establish and maintain DNA methylation at these three sequence contexts, respectively. CHROMOMETHYLASES (CMTs) are an important class of plant-specific DNA methylation enzymes, which are characterized by the presence of a CHRromatin Organisation MOdifier (CHROMO) domain between the cytosine methyltransferase catalytic motifs I and IV [1]. Identification, expression, and functional characterization of CMTs have been extensively performed in the model plant *Arabidopsis thaliana* [2, 3, 4] and in the model grass species *Zea mays* [5, 6, 7].

There are three CMT genes encoded in the *A. thaliana* genome: CMT1, CMT2, and CMT3 [2, 8, 9, 10]. CMT1 is the least studied of the three CMTs as a handful of *A. thaliana* accessions contain an Evelknievel retroelement insertion or a frameshift mutation truncating the protein, which suggested that CMT1 is nonessential [8]. The majority of DNA methylation at CHH sites (mCHH) at long transposable elements in pericentromeric regions of the genome is targeted by a CMT2-dependent pathway [3, 4]. Allelic variation at CMT2 has been shown to alter genome-wide levels of CHH DNA methylation (mCHH), and plastic alleles of CMT2 may play a role in adaptation to temperature [11, 12, 13]. In contrast, DNA methylation at CHG (mCHG) sites is often maintained by CMT3 through a reinforcing loop with histone H3 lysine 9 di-methylation (H3K9me2) catalyzed by the KRYPTONITE (KYP)/SU(VAR)3-9 HOMOLOG 4 (SUVH4), SUVH5 and SUVH6 lysine methyltransferases [2, 6, 14, 15]. In *Z. mays*, ZMET2 is a functional homolog of CMT3 and catalyzes the maintenance of mCHG [5, 6, 7]. A paralog of ZMET2, ZMET5, contributes to the maintenance of mCHG to a lesser degree in *Z. mays* [5, 7]. Homologous CMTs have been identified in other flowering plants (angiosperms) [16, 17, 18, 19]: the moss *Physcomitrella patens*, the lycophyte *Selaginella moellendorffii*, and the green algae *Chlorella sp.* NC64A and *Volvox carteri* [20]. The function of CMTs in species sister to angiosperms (flowering plants) is poorly understood. However, in at least *P. patens* a CMT protein contributes to mCHG [21].

A large number of genes in angiosperms exclusively contain CG DNA methylation (mCG) in the transcribed region and a depletion of mCG from both the transcriptional start and stop sites (referred to as “*gene body DNA methylation*”, or “*gbM*”) [22, 23, 24, 25]. GbM genes are generally constitutively expressed, evolutionarily conserved, and typically longer than un-methylated genes [25, 26, 27]. How gbM is established and subsequently maintained is unclear. However, recently it was discovered that CMT3 has been independently lost in two angiosperm species belonging to the Brassicaceae family of plants and this coincides with the loss of gbM [19, 25]. Furthermore, *A. thaliana* and closely related Brassicaceae species have reduced levels of mCHG on a per cytosine basis, but still posses CMT3 [19, 25], which indicates changes at the molecular level may have disrupted function of CMT3. This has led to a hypothesis that the evolution of gbM is linked to incorporation/methylation of histone H3 lysine-9 di-methylation (H3K9me2) in gene bodies with subsequent failure of INCREASED IN BONSAI METHYLATION 1 (IBM1) to de-methylate H3K9me2 [19, 28]. This provides a substrate for CMT3 to bind and methylate DNA, and through an unknown mechanism leads to mCG. MCG is maintained over evolutionary timescales by the CMT3-dependent mechanism and during DNA replication by the maintenance DNA METHYTRANSFERASE 1 (MET1). Methylated DNA then provides a substrate for binding by KRYPTONITE (KYP) and related family members through their SRA domains, which increases the rate at which H3K9 is di-methylated [29]. Finally, mCG spreads throughout the gene over evolutionary time [19]. A similar model was previously proposed, which links gene body mCG with transcription, mCHG, and IBM1 activity [28].

Previous phylogenetic studies have proposed that CMT1 and CMT3 are more closely related to each other than to CMT2, and that ZMET2 and ZMET5 proteins are more closely related to CMT3 than to CMT1 or CMT2 [5], and the placement of non-seed plant CMTs more closely related to CMT3 [21]. However, these studies were not focused on resolving phylogenetic relationships within the CMT gene family, but rather relationships of CMTs between a handful of species. These studies have without question laid the groundwork to understand CMT-dependent DNA methylation pathways and patterns in plants. However, the massive increase in transcriptome data from a broad sampling of plant species together with advancements in sequence alignment and phylogenetic inference algorithms have made it possible to incorporate thousands of sequences into a single phylogeny, allowing for a more complete understanding of the CMT gene family. Understanding the evolutionary relationships of CMT proteins is foundational for inferring the evolutionary origins, maintenance, and consequences of genome-wide DNA methylation and gbM.

Here we investigate phylogenetic relationships of CMTs at a much larger evolutionary timescale using data generate from the 1KP Consortium (www.onekp.com). In the present study we have analyzed 771 mRNA transcripts and annotated genomes, identified as belonging to the CMT gene family, from an extensive taxonomic sampling of 443 different species including eudicots (basal, core, rosid, and asterid), basal angiosperms, monocots and monocots/commelinid, magnoliids, gymnosperms (conifers, Cycadales, Ginkgoales), monilophytes (ferns and fern allies), lycophytes, bryophytes (mosses, liverworts and hornworts), and green algae. CMT homologs identified across Viridiplantae (land plants and green algae) indicate that CMT genes originated prior to the origin of Embryophyta (land plants) (≥480 MYA) [30, 31, 32, 33]. In addition, phylogenetic relationships suggests at least two duplication events occurred within the angiosperm lineage giving rise to the CMT1, CMT2, and CMT3 gene clades. In the light of CMT evolution we explored patterns of genomic and genic DNA methylation levels in 77 species of Viridiplantae, revealing diversity of the epigenome within and between major taxonomic groups, and the evolution of gbM in association with the origin of CMT3 and orthologous sequences.

## RESULTS

### The origins of CHROMOMETHYLASES

CMTs are found in most major taxonomic groups of land plants and some algae: eudicots, basal angiosperms, monocots and commelinids, magnoliids, gymnosperms, ferns, lycophytes, mosses, liverworts, hornworts, and green algae (Fig. 1a and Table S1). CMTs were not identified in transcriptome data sets for species sister to Viridiplantae including those belonging to Glaucophyta, red algae, Dinophyceae, Chromista, and Euglenozoa. CMTs were identified in a few green algae species: *Picocystis salinarum*, *Cylindrocystis sp*., and *Cylindrocystis brebissonii*. Additionally, functional CMTs – based on presence/absence of characterized protein domains – were not identified from three species within the gymnosperm order Gnetales. A transcript with a CHROMO and C-5 cytosine-specific DNA methylase domain was identified in *Welwitschia mirabilis* (Gnetales), but this transcript did not include a Bromo Adjacent Homology (BAH) domain. The BAH domain is an interaction surface that is required to capture H3K9me2, and mutations that abolish this interaction causes a failure of a CMT protein (i.e., ZMET2) binding to nucleosomes and a complete loss of activity *in vivo* [6]. Therefore, although a partial sequence is present, it might represent a nonfunctional allele of a CMT. Alternatively, it might represent an incomplete transcript generated during sequencing and assembling of the transcriptome. Overall, the presence of CMT homologs across Viridiplantae and their absence from sister taxonomic groups suggest CMT evolved following the divergence of green algae [35, 35].

**Fig. 1.**
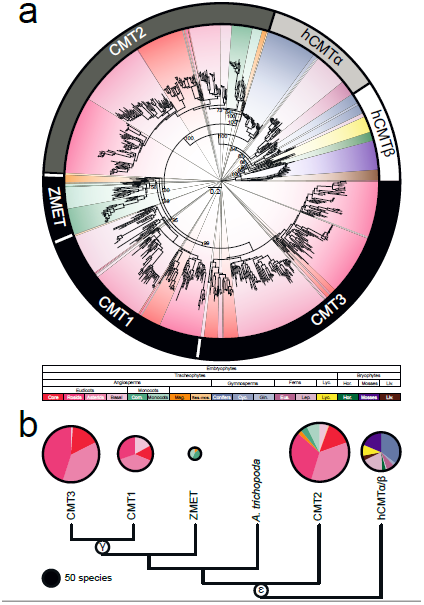
Phylogenetic relationships of CMTs across Embryophyta. **a**, CMT sare separated into four monophyletic clades based on bootstrap support and the relationship of *A. thaliana* CMTs: (i) the gbM-dependent CMT superclade with subclades CMT1, CMT3, ZMET and *A. trichopoda*; (ii) CMT2 and; (iii) homologous (hCMT) α and β. CMT1 and CMT3 clades only contain eudicot species of plants suggesting a eudicot-specific duplication event that occurred after the divergence of eudicots from monocots and monocots/commelinids. Sister to CMT1 and CMT3 is the monophyletic group ZMET, which contains monocots, monocots/commelinids, and magnoliids. CMT2 is sister to CMT1 and CMT3. Lastly, the polyphyletic hCMT clades are sister to all previously mentioned clades. HCMTα is sister to CMT2 and the CMT superclade and contains gymnosperm and ferns. HCMTβ contains gymnosperms, ferns and other early diverging land plants. **b**, A collapsed CMT gene family tree showing the seven clades described in **a**. Pie charts represent species diversity within each clade, and are scaled to the number of species. Two duplication events shared by all angiosperms (ε) and eudicots () gave rise to what is now referred to as CMT1, CMT2 and CMT3. These duplication events correspond to what was reported by Jiao et al. (2011). Values at nodes in **a** and **b** represent bootstrap support from 1000 replicates, and **a** was rooted to the clade containing all liverwort species.

The relationships among CMTs suggest that CMT2 and the clade containing CMT1, CMT3 and ZMET arose from a duplication event at the base of all angiosperms (Fig. 1b). This duplication event might have coincided with event ε, the ancestral angiosperm whole genome duplication (WGD) event [36]. Relationships among clades sister to angiosperm CMTs largely recapitulate species relationships (Fig. 1a) [34, 37]. However, CMTs in gymnosperms and ferns are paraphyletic (Fig. 1a). Similarly, these homologous sequences might have been derived from a WGD (i.e., ζ, the ancestral seed plant WGD), with one paralog being the ancestor to CMT1, CMT2 and CMT3, and ZMET [36]. Previously identified CMTs in *S. moellendorffii* [20] and *P. patens* [20, 38] were identified, which are sister to clades containing CMT1, CMT2, and CMT3 and ZMET (Fig. 1a). CMTs previously identified in the green algae *Chlamydomonas reinhardtii*, *Chlorella sp.* NC64A and *Volvox carteri* were excluded from phylogenetic analysis because they lacked the CHROMO and other domains typically associated with CMT proteins (Figure S1). Furthermore, based on percent amino acid identity *C. reinhardtii* and *V. carteri* CMT sequences are homologous to MET1 (Table S2). Similar to *S. moellendorffii* and *P. patens* CMT sequences, green algae CMT sequences are sister to clades containing CMT1, CMT2, and CMT3 and ZMET (Figure S2). The increase taxonomic sampling redefines relationships of CMTs in early-diverged land plants and in Viridiplantae in general [18, 20, 21, 38, 39].

Further diversification of CMT proteins occurred in eudicots. CMT1 and CMT3 clades contain only sequences from eudicots (Fig. 1a and b). This relationship supports the hypothesis that CMT1 and CMT3 arose from a duplication event shared by all eudicots. Thus, CMT1 and CMT3 might be the result of the γ WGD event at the base of eudicots [36]. Synteny between CMT1 and CMT3, despite ~125 million years of divergence, further supports this hypothesis (Figure S3a). Not all eudicots possess CMT1, CMT2, and CMT3, but rather exhibit CMT gene content ranging from zero to three (Figure S4a). Also, many species possess multiple copies of CMT1, CMT2, or CMT3. The presence/absence of CMTs might represent differences in transcriptome sequencing coverage or spatial and temporal divergence of expression. However, CMT2, CMT3, and homologous proteins have functions in methylating a significant number of non-CG sites throughout the entire genome and thus are broadly expressed in *A. thaliana*, *Z. mays* and other species [8, 6, 18, 25]. Additionally, eudicot species with sequenced and assembled genomes show variation in the presence/absence and copy number of CMTs. Hence the type of tissue(s) used in transcriptome sequencing (www.onekp.com) would have limited biases against CMTs, suggesting that the variation reflects presence/absence at a genetic level.

The *Z. mays* in-paralogs ZMET2 and ZMET5, and closely related CMTs in other monocots, commelinids, and magnoliids form a well-supported monophyletic clade (Fig. 1a and Figure S3b and c). In addition to *Z. mays*, in-paralogs were identified in *Sorghum bicolor* and *Brachypodium distachyon* (Fig. 1a and Figure S3b). Relationships of *S. bicolor* and *Z. mays* ZMETs differed between gene and amino acid derived phylogenies (Fig. 1a and Figure S3b). However, synteny between paralogs of both species supports two independent duplications (Figure S3c). Also, paralogous ZMETs are shared across species (Figure S3b). These shared paralogs might have originated from a Poaceae-specific duplication event, which was followed by losses in some species. The contribution of each paralog to DNA methylation and other chromatin modifications remains unknown at this time.

Akin to eudicots, monocots and monocots/commelinids possess combinations of ZMET and CMT2 (Figure S4b). For example, the model grass species *Z. mays* has lost CMT2, whereas the closely related species *S. bicolor* possess both ZMET and CMT2 (Table S1). ZMET is not strictly homologous to CMT3, and represents a unique monophyletic group that is sister to both CMT1 and CMT3. However, ZMET2 is functionally homologous to CMT3 and maintains DNA methylation at CHG sites [6, 7]. Unlike CMT3, ZMET2 is associated with DNA methylation at CHH sites within some loci [7]. Given the inclusion of monocot and magnoliid species in the monophyletic ZMET clade, this dual-function is expected to be present in other monocot species, and in magnoliid species.

Overall, these redefined CMT clades, and monophyletic clades of broad taxonomic groups, are well supported (Fig. 1a). Thus, the identification of novel CMTs in magnoliids, gymnosperms, lycophytes, hornworts, liverworts, bryophytes, and green algae pushes the timing of evolution of CMT, and potentially certain mechanisms maintaining mCHG and mCHH, prior to the origin of Embryophyta (≥480 million years ago [MYA]) [30, 31, 32, 33].

### Reduced selective constraint of CMT3 in the Brassicaceae affects gbM

Recent work has described the DNA methylomes of 34 angiosperms, revealing extensive variation across this group of plants [25]. This variation was characterized in terms of levels of DNA methylation, and number of DNA methylated genes. DNA methylation levels describe variation within a population of cells. Although ascribing genes as DNA methylated relies on levels of DNA methylation, this metric provides insights into the predominant DNA methylation pathway and expected relationship to genic characteristics [25, 26, 27]. The genetic underpinnings of this variation are not well understood, but some light has been shed through investigating DNA methylation within the Brassicaceae [19]. The Brassicaceae have reduced levels of genomic and genic levels of mCG, genome-wide per-site levels of mCHG, and numbers of gbM genes [19, 25]. In at least *E. salsugineum* and *C. planisiliqua* this reduction in levels of DNA methylation and numbers of gbM genes has been attributed to the loss of the CMT3 [19]. However, closely related species with CMT3 – *Brassica oleracea*, *Brassica rapa* and *Schrenkiella parvula* – have reduced levels of gbM and numbers of gbM genes compared to the sister clade of *A. thaliana*, *Arabidopsis lyrata*, and *Capsella rubella* (Fig. 2a and b) and overall to other eudicots [19, 25]. Although CMT3 is present, changes at the sequence level, including the evolution of deleterious or functionally null alleles, could disrupt function to varying degrees. At the sequence level, CMT3 has evolved at a higher rate of molecular evolution – measured as *dN/dS* (ω) – in the Brassicaceae (ω=0.175) compared to 162 eudicots (ω=0.097), with further increases in the clade containing *B. oleracea*, *B. rapa* and *S. parvula* (ω=0.241) compared to the clade containing *A. thaliana*, *A. lyrata* and *C. rubella* (ω=0.164) (Fig. 2). A low background rate of molecular evolution suggests purifying selection acting to maintain low allelic variation across eudicots. Conversely, increased rates of molecular evolution can be a consequence of positive selection. However, a hypothesis of positive selection was not preferred to contribute to the increased rates of ω in either Brassicaceae clade (Table S3). Alternatively, relaxed selective constraint possibly resulted in an increased ω, which might have introduced null alleles ultimately affecting function of CMT3, and in turn, affecting levels of DNA methylation and numbers of gbM genes. The consequence of the higher rates of molecular evolution in the clade containing *Brassica spp*. and *S. parvula* relative to all eudicots and other Brassicaceae are correlated with an exacerbated reduction in the numbers of gbM loci and their methylation levels, which suggests unique substitutions between clades or a quantitative affect to an increase in the number of substitutions. However, at least some substitutions affecting function are shared between Brassicaceae clades because both have reductions in per-site levels of mCHG [19].

**Fig. 2.**
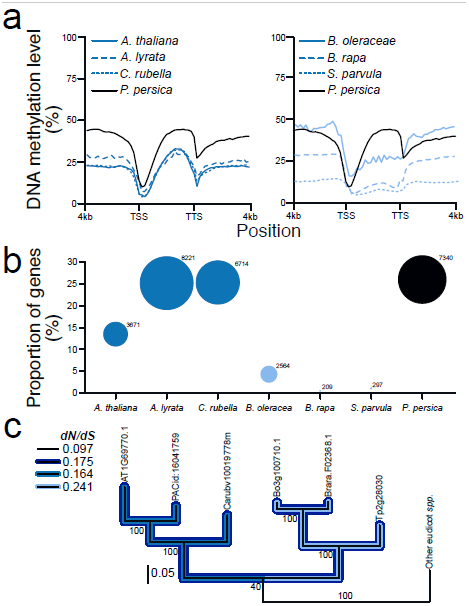
Non-neutral evolution of CMT3 in the Brassicaceae is correlated with reduced levels of genic mCG and numbers of gbM loci. **a**, Distribution of mCG upstream, downstream and within gene bodies of Brassicaceae species and outgroup species *Prunus persica*. MCG levels within gene bodies of Brassicaceae species are within the bottom 38% of 34 angiosperms. Data used represents a subset of that previously published by [19] and [25]. TSS: transcriptional start site; and TTS: transcriptional termination site. **b**, Similarly the number of gbM genes within the genome of Brassicaceae species are within the bottom 15% of 34 angiosperms. The size of the circle corresponds to the number of gbM genes within each genome. Data used represents a subset of that previously published by [19] and [25]. **c**, Changes at the amino acid level of CMT3 is correlated to reduced genic levels of DNA methylation and number of gbM genes in the Brassicaceae. An overall higher rate of molecular evolution measured as the number of non-synonymous substitutions per non-synonymous site divided by the number of synonymous substitutions per synonymous site (ω) was detected in the Brassicaceae. Also, a higher rate ratio of ω was detected in the Brassicaceae clade containing *B. rapa* and closely related species compared to the clade containing *A. thaliana* and closely related species. The higher rate ratio in the Brassicaceae, compared the background branches, was not attributed to positive selection.

### Divergence of DNA methylation patterns within gene bodies during Viridiplantae evolution

Levels and distributions of DNA methylation within gene bodies are variable across Viridiplantae. Levels of mCG range from ~2% in *S. moellendorffi* to ~86% in *Chlorella sp*. NC64A (Fig. 3a). Other plant species fall between these two extremes (Fig. 4a) [25]. *Beta vulgaris* remains distinct among angiosperms and Viridiplantae with respect to levels of DNA methylation at all sequence contexts (Fig. 3a). Similarly, *Z. mays* is distinct among monocots and monocots/commelinids (Fig. 3a). Gymnosperms and ferns possess similar levels of mCG to mCHG within gene bodies and levels of mCHG qualitatively parallel those of mCG similar to observations in recently published study (Fig. 3a and Figure S5) [40]. A similar pattern is observed in *Z. mays*. However, this pattern is not shared by other monocots/commelinids [25]. High levels of mCHG is common across the gymnosperms and ferns investigated in this study, and tends to be higher than levels observed in angiosperms (Fig. 3a and Figure S5) [25]. DNA methylation at CG, CHG and CHH sites within gene bodies was detected in the liverwort *Marchantia polymorpha*) (Fig. 3a). Furthermore, DNA methylation at CG sites was not detected in the *P. patens* when all genes are considered (Fig. 3a). Overall, increased taxonomic sampling has revealed natural variation between and within groups of Viridiplantae.

**Fig. 3.**
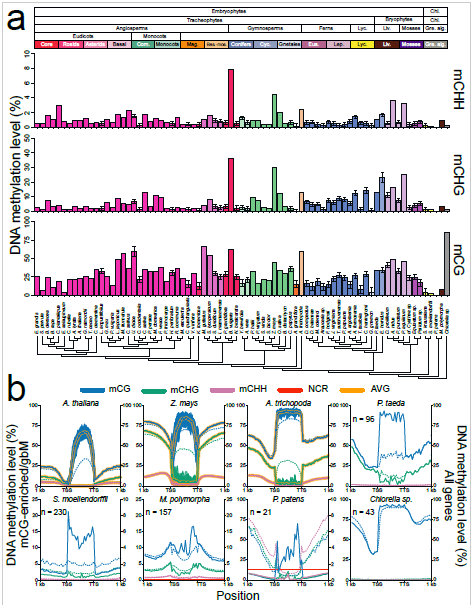
Variation in levels of DNA methylation within gene bodies across 668 Viridiplantae. **a**, DNA methylation at CG, CHG, and CHH sites within gene bodies can be found at the majority of species investigated. Variation of DNA methylation levels within gene bodies at all sequence contexts is high across all land plants, and within major taxonomic groups. mCG levels are typically higher than mCHG, followed by mCHH. However, levels of mCG and mCHG within genes are similar in gymnosperms and ferns. Error bars represent 95% confidence intervals for species with low sequencing coverage. Cladogram was generated from Open Tree of Life [53]. **b**, The distribution of DNA methylation within genes (all [dashed lines] and mCG-enriched/gbM [solid lines]) has diverged among taxonomic groups of Viridiplantae represented by specific species. Based on the distribution of DNA methylation, and number of mCG-enriched genes, gbM is specific to angiosperms. However, mCG-enriched genes in *P. taeda* share some DNA methylation characteristics to *A. trichopoda*. However, other characteristics associated with gbM genes remains unknown at this time for mCG-enriched genes in gymnosperms and other early diverging Viridiplantae. The yellow-highlighted line represents the average from 100 random sampling of 100 gbM genes in angiosperms and was used to assess biases in numbers of mCG-enriched genes identified. NCR: non-conversion rate; TSS: transcriptional start site; and TTS: transcriptional termination site.

**Fig. 4.**
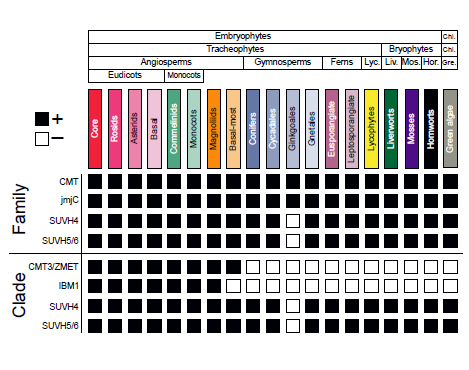
Presence/absence (+/−) of genes likely involved in the evolution of gbM and heterochromatin formation for various taxonomic groups of Viridiplantae. Families (orthogroups) of gbM- and heterochromatin-related genes are taxonomically diverse. However, after phylogenetic resolution, clades containing proteins of known function in *A. thaliana* are less diverse. Specifically, the CMT3 and orthologous genes (ZMET2 and ZMET5, and *A. trichopoda* CMT3), and IBM1 are angiosperm-specific. Other clades – SUVH4 and homologous SUVH5/6 (hSUVH5/6) – are more taxonomically diverse, which might relate to universal functions in heterochromatin formation.

Despite the presence of mCG within the gene bodies of angiosperms, gymnosperms, ferns, lycophytes, liverworts, and green algae; the distributions across gene bodies differ (Fig. 3b). In angiosperms (eudicots, commelinids, monocots and basal angiosperms) CG DNA methylation is depleted at the transcriptional start and termination sites (TSS and TTS, respectively), and gradually increases towards the center of the gene body (Fig. 3b). In the basal angiosperm *Amborella trichopoda*, levels of mCG decline sharply prior to the TTS (Fig. 3b). Similar to angiosperms, mCG is reduced at the TSS relative to the gene body in the gymnosperm *Pinus taeda* (Fig. 4b). However, mCG is not reduced at the TTS (Fig. 3b). Additionally, DNA methylation at non-CG (mCHG and mCHH) sites is not reduced at the TSS and TTS. Little difference in mCG, mCHG, and mCHH within gene bodies, and upstream and downstream regions are observed in *S. moellendorffii* (Fig. 3b). Additionally, mCG, mCHG, and mCHH are not excluded from the TSS and TTS (Fig. 3b). As opposed to angiosperms and gymnosperms (*P. taeda*), mCG in *M. polymorpha* decreases towards the center of the gene body (Fig. 3b). This distribution also occurs for methylation at non-CG sites in *M. polymorpha* (Fig. 3b). Additionally, *M. polymorpha* has distinctive high levels of mCG, mCHG, and mCHH surrounding the TSS and TTS (Fig. 3b). Finally, in *Chlorella sp*. NC64A, mCG is enriched at near 100% across the entire gene body (Fig. 3b).

The presence of mCG within gene bodies indicates that a gene could possess gbM. However, other types of DNA methylated genes contain high levels of mCG [25], thus enrichment tests were performed to identify genes that are significantly enriched for mCG and depleted of non-CG methylation (i.e., gbM genes). Genes matching this DNA methylation enrichment profile were identified in species sister to angiosperms: gymnosperms, lycophytes, liverworts, mosses and green algae (Fig. 3b). The proportion of genes within each genome or subset of the genome (*P. taeda*) was small compared to angiosperms with gbM (Fig. 3b). Furthermore, the number of gbM genes was comparable to angiosperms without gbM, which suggests these identified genes are the result of statistical noise (Figure S6a). This is most likely the case for lycophytes, liverworts, mosses and green algae, since the levels of mCG within genes bodies is highly skewed (Figure S7). Additionally, the distribution of mCG and non-CG methylation across the gene body is unlike the distribution of gbM genes (Fig. 3b) [25]. However, the gymnosperm *P. taeda* shares some similarities to gbM genes of the basal angiosperm *A. trichopoda* (Fig. 3b). Hence, mCG-enriched genes identified in gymnosperms, lycophytes, liverworts, mosses and green algae are most likely not gbM.

### Correlated evolution of CMT3 and the histone de-methylase IBM1 in angiosperms

The exact mechanisms by which genes are targeted to become gbM and the establishment of DNA methylation at CG sites is currently unknown. One proposed possibility is the failure of IBM1 to remove H3K9me2 within genes. This would provide the necessary substrate for CMT3 to associate with nucleosomes in genes. Due to the tight association between CMT3 and IBM1 (and SUVH4/5/6) these proteins might have evolved together. Resolution of phylogenetic relationships supports monophyly of IBM1 and orthologous sequences that is unique to angiosperms (Fig. 4 and Figure S8). Furthermore, high levels of mCHG and/or similar levels of mCHG to mCG in gymnosperms, ferns, *S. moellendorffii* (lycophyte), *M. polymorpha* (liverwort) and *P. patens* (moss) compared to angiosperms suggests a functionally homologous histone de-methylase is not present in these taxonomic groups and species. The absence of IBM1 in the basal-most angiosperm *A. trichopoda* and similarities of DNA methylation distribution between gbM genes and *P. taeda* mCG-enriched genes further supports a role of CMT3 and IBM1 in maintenance of mCG within gene bodies. Unlike CMT3 and IBM1, histone methylases SUVH4 and SUVH5/6 are common to all taxonomic groups investigated, which suggests common ancestry and shared functions of transposon silencing (Fig. 4 and Figures S8 and S9) [22, 41, 42, 43]. However, a Brassicaceae-specific duplication event gave rise to SUVH5 and SUVH6, and other Viridiplantae possess a homologous SUVH5/6 (hSUVH5/6) (Figure S9b and c). Additionally, a duplication event shared by all monocots and monocots/commelinids generated paralogous hSUVH5/6, and additionally duplication event occurred in the Poaceae (Figure S9d). The duplication event that gave rise to ZMET paralogs in the Poaceae might have also generated the paralogous hSUVH5/6. The diversity in levels and patterns of DNA methylation within gene bodies suggests corresponding changes in function of DNA methyltransferases and/or histone de-methylases during Viridiplantae divergence. Furthermore, monophyletic, angiosperm-specific clades of a gbM-dependent CMT and IBM1 suggest co-evolution of proteins involved in the gbM pathway.

## DISCUSSION

CMTs are conserved DNA methyltransferases across Viridiplantae. Evolutionary phenomenon and forces have shaped the relationships of CMTs, which have most likely contributed to functional divergence among and within taxonomic groups of Viridiplantae. Duplication events have contributed to the unique relationships of CMTs, and have given rise to clade-, family- and species-specific CMTs. This includes the eudicot-specific CMT1 and CMT3, paralogous CMTs within monocots/commelinids (ZMETs), and the *Z. mays*-specific ZMET2 and ZMET5. The paralogous CMT1 and CMT3, and ortholgous CMTs in monocots, monocots/commelinids, magnoliids and basal angiosperms form a superclade that is sister to CMT2. Homologous CMTs in gymnosperms and ferns are paraphyletic, and clades are sister to all CMTs – including CMT1, CMT2, CMT3 and ZMET – in angiosperms. CMTs have been shown to maintain methylation at CHG sites (CMT3 and ZMET5, and hCMTβ in *P. patens*) and methylate CHH sites within deep heterochromatin (CMT2) [2, 4, 6, 11, 14, 15, 20, 21], whereas CMT1 is nonfunctional in at least *A. thaliana* accessions [8]. However, recent work has provided evidence for the role of CMT3 in the evolution of mCG within gene bodies, and specifically gbM, within angiosperms [19]. Additionally, non-neutral evolution of CMT3 can affect levels of genome-wide mCHG and within gene body mCG, and the number of gbM genes. Hence, functional divergence following duplication might be more widespread [44], and the exact fate of paralogous CMTs and interplay between paralogs in shaping the epigenome remain unknown at this time.

DNA methylation within genes is common in Viridiplantae. However, certain classes of DNA methylated genes might be specific to certain taxonomic groups within the Viridiplantae. GbM is a functionally enigmatic class of DNA methylated genes, which is characterized by an enrichment of mCG and depletion of non-mCG within transcribed regions and depletion of DNA methylation from all sequence contexts at the TSS and TTS. These genes are typically constitutively expressed, evolutionary conserved, housekeeping genes, which compose a distinct proportion of protein coding genes [19, 25, 26, 27, 45]. GbM genes have been mostly studied in angiosperms and evidence for the existence of this class of DNA methylated gene outside of angiosperms is limited [40]. However, in the present study, genes matching the DNA methylation profile of gbM genes – enrichment of mCG and depletion of non-mCG – were identified in taxonomic groups sister to angiosperms: gymnosperms, lycophytes, liverworts, mosses and green algae. It is unclear if these genes are gbM genes in light of findings in angiosperms [19, 25, 26, 27]. For example, the low proportion of mCG-enriched genes supports the absence of gbM in gymnosperms, lycophytes, liverworts, mosses and green algae. Additionally, the distribution of mCG among all genes and across the gene body of mCG-enriched genes supports the absence of gbM in lycophytes, liverworts, mosses and green algae. However, similar distributions of mCG between gbM genes in the basal angiosperm *A. trichopoda* and mCG-enriched genes in the gymnosperm *P. taeda* are observed, which support the presence of gbM in this species and possibly other gymnosperms. Also, a small proportion of mCG-enriched genes in gymnosperms are homologous to gbM genes in *A. thaliana* (Figure S6b). With that being said, sequence conservation is not the most robust indicator of gbM [25]. GbM genes compose a unique class of genes with predictable characteristics [19, 25, 26, 27]. Through comparison of mCG-enriched genes identified in early diverging Viridiplantae to angiosperms with and without gbM, there is stronger support that this epigenetic feature is unique to angiosperms. However, future work including deeper WGBS and RNA-seq, and additional and improved genome assemblies – especially for gymnosperms and ferns – will undoubtedly contribute to our understanding of the evolution of gbM.

GbM is dependent on the CHG maintenance methyltransferase CMT3 or an orthologous CMT in angiosperms. Support for the dependency of gbM on CMT3 comes from the naturally occurring Δ*cmt3* mutants *E. salsugineum* and *C. planisilqua*, which is correlated with the lack of gbM genes [19, 25]. The independent loss of CMT3 has also affected mCHG with low overall and per-site levels recorded for these species [25]. Both species belong to the Brassicaceae family, and other species within this family show reduced numbers of gbM genes compared to other eudicot and angiosperm species [25]. Although CMT3 is present in these species, relaxed selective constraint might have introduced alleles which functionally compromise CMT3 resulting in decreased per-site levels of mCHG and the number of gbM genes [25]. The functional compromises of CMT3 non-neutral evolution are shared and have diverged between clades of Brassicaceae, respectively, which might reflect shared ancestry between clades and the unique evolutionary history of each clade. Furthermore, more relaxed selective constraint – as in the *Brassica spp*. and *S. parvula* – is correlated with a more severe phenotype relative to the other Brassicaceae clade. The dependency of gbM on a CMT protein might extend into other taxonomic groups of plants. Phylogenetic relationships of CMTs found in Viridiplantae and the location of *A. thaliana* CMTs support a eudicot-specific, monophyletic CMT3 clade. The CMT3 clade is part of a superclade, which includes a monophyletic clade of monocot (ZMET) and magnoliid CMTs, and a CMT from the basal angiosperm *A. trichopoda*. Thus, the CMT-dependent gbM pathway might be specific to angiosperms. However, a homologous, closely related CMT in gymnosperms and ferns (i.e., hCMTα) might have a similar function. It is conceivable that other proteins and chromatin modifications that interact with CMTs and non-CG methylation are important for the evolution of gbM, and thus have evolved together. Specifically, IBM1 that de-methylates H3K9me2 and SUVH4/5/6 that binds to H3K9me2 and methylates CHG sites both act upstream of CMT3. One proposed model for the evolution of gbM requires failure of IBM1 and rare mis-incorporation of H3K9me2, which initiates mCHG by SUVH4/5/6 and maintenance by CMT3 [19, 28]. IBM1 shares similar patterns and taxonomic diversity as CMT3 and orthologous CMTs involved in gbM. Also, unlike most angiosperms investigated to date – with *A. trich*opoda as the exception – gymnosperms and ferns do not possess an IBM1 ortholog, hence IBM1 might be important for the distribution of mCG within gene bodies. Furthermore, the lack of IBM1 in *A. trichopoda* and *P. taeda* might explain some similarities shared between gbM genes and mCG-enriched genes with respect to the deposition of mCG, respectively. However, the exact relationship between gbM and IBM1 is unknown and similarities in underlying nucleotide composition of genes might affect distribution of mCG. Overall, the patterns of DNA methylation within gene bodies and the phylogenetic relationships of CMTs support a CMT3 and orthologous CMT-dependent mechanism for the maintenance of gbM in angiosperms, which is stochastically initiated by IBM1.

## CONCLUSIONS

In summary, we present the most comprehensive CMT gene-family phylogeny to date. CMTs are ancient proteins that evolved prior to the diversification of Embryophyta. A shared function of CMTs is the maintenance of DNA methylation at non-CG sites, which has been essential for DNA methylation at long transposable elements in the pericentromeric regions of the genome [6, 14, 15]. However, CMTs in some species of eudicots have been shown to be important for mCG within gbM genes [19]. Refined relationships between CMT1, CMT2, CMT3, ZMET, and other homologous CMT clades have shed light on current models for the evolution of gbM, and provided a framework for further research on the role of CMTs in establishment and maintenance of DNA methylation and histone modifications. Patterns of DNA methylation within gene bodies have diverged between Viridiplantae. Other taxonomic groups do not share the pattern of mCG associated with gbM genes in the majority of angiosperms, which further supports specificity of gbM in angiosperms. However, genic DNA methylation commonalities between angiosperms and other taxonomic groups were identified. DNA methylation within gene bodies and its consequences of or relationship to expression and other genic features has been extensively studied in angiosperms [25] and shifting focus to other taxonomic groups of plants for deep methylome analyses will aid in understanding the shared consequences of genic DNA methylation. Understanding the evolution of additional chromatin modifiers will undoubtedly unravel the epigenome and reveal unique undiscovered mechanisms.

## METHODS

### 1KP sequencing, transcriptome assembling and orthogrouping

The One Thousand Plants (1KP) Consortium includes assembled transcriptomes and predicted protein coding sequences from a total of 1329 species of plants (Table S1). Additionally, gene annotations from 24 additional species – *Arabidopsis lyrata, Brachypodium distachyon, Brassica oleracea, Brassica rapa, Citrus clementina, Capsella rubella, Cannabis sativa, Cucumis sativus, Eutrema salsugineum, Fragaria vesca, Glycine max, Gossypium raimondii, Lotus japonicus, Malus domestica, Marchantia polymorpha, Medicago truncatula, Panicum hallii, Panicum virgatum, Pinus taeda, Physcomitrella patens, Ricinus communis, Setaria viridis, Selaginella moellendorffii*, and *Zea mays* – were included (https://phytozome.jgi.doe.gov/pz/portal.html and http://pinegenome.org/pinerefseq/). The CMT gene family was extracted from the previously compiled 1KP orthogroupings using the *A. thaliana* gene identifier for CMT1, CMT2 and CMT3. A single orthogroup determined by the 1KP Consortium included all three *A. thaliana* CMT proteins, and a total of 5383 sequences. Sequences from species downloaded from Phytozome, that were not included in sequences generated by 1KP, were included to the gene family through reciprocal best BLAST with *A. thaliana* CMT1, CMT2 and CMT3. In total the CMT gene family included 5449 sequences from 1043 species. We used the protein structure of *A. thaliana* as a reference to filter the sequences found within the CMT gene family. Sequences were retained if they included the same base PFAM domains as *A. thaliana* – CHROMO, BAH, and C-5 cytosine-specific DNA methylase domains – as identified by Interproscan [46]. These filtered sequences represent a set of high-confident, functional, ideal CMT proteins, which included 771 sequences from 432 species, and were used for phylogenetic analyses.

### Phylogeny construction

To estimate the gene tree for the CMT sequences, a series of alignment and phylogenetic estimation steps were conducted. An initial protein alignment was carried out using Pasta with the default settings [47]. The resulting alignment was back-translated using the coding sequence (CDS) into an in-frame codon alignment. A phylogeny was estimated by RAxML [48] (-m GTRGAMMA) with 1000 rapid bootstrap replicates using the in-frame alignment, and with only the first and second codon positions. Long branches can effect parameter estimation for the substitution model, which can in turn degrade phylogenetic signal. Therefore, phylogenies were constructed with and without green algae species, and were rooted to the green algae clade or liverworts, respectively. The species *Balanophora fungosa* has been reported to have a high substitution rate, which can also produce long branches, and was removed prior to phylogenetic analyses. Identical workflows were used for jumonji (jmjC) domain-containing (i.e., IBM1), SUVH4, and SUVH5/6 gene families.

### Codon analysis

Similar methodology as described above was used to construct phylogenetic trees for testing hypotheses on the rates of evolution in a phylogenetic context. However, the program Gblocks [49] was used to identify conserved codons. The parameters for Gblocks were kept at the default settings, except allowing for 50% gapped positions. The program Phylogenetic Analysis by Maximum Likelihood (PAML) [50] was used to test branches (branch test) and sites along branches (branch-site test) for deviations from the background rate of molecular evolution (ω) and for deviations from the neutral expectation, respectively. Branches tested and a summary of each test can be found in Table S3.

### MethylC-seq

MethylC-seq libraries were prepared according to the following protocol [51]. For *A. thaliana, A. trichopoda, Chlorella sp*. NC64A, *M.polymorpha, P. patens, P. taeda, S. moellendorffii*, and *Z. mays* reads were mapped to the respective genome assemblies. *P. taeda* has a large genome assembly of ~23 Gbp divided among ~14k scaffolds (http://dendrome.ucdavis.edu/ftp/Genome_Data/genome/pinerefseq/Pita/v1.01/README.txt). Due to computational limitations imposed by the large genome size only 4 Gbp of the *P. taeda* genome assembly was used for mapping, which includes 2411 (27%) of the high quality gene models. Prior to mapping for species with only transcriptomes each transcript was searched for the longest open reading frame from all six possible frames, and only transcripts beginning with a start codon and ending with one of the three stop codons were kept. All sequencing data for each species was aligned to their respective transcriptome or species within the same genus using the methylpy pipeline [52]. All MethylC-seq data used in this study can be found in Tables S4 and S5. Weighted methylation was calculated for each sequence context (CG, CHG and CHH) by dividing the total number of aligned methylated reads by the total number of methylated plus un-methylated reads. Since, per site sequencing coverage was low – on average ~1× – subsequent binomial tests could not be performed for the majority of species to bin genes as gbM [25]. To investigate the affect of low coverage we compared levels of DNA methylation of 1× randomly sampled MethylC-seq reads to actual levels for 32 angiosperm species, *S. moellendorffii* (lycophyte), *M. polymorpha* (liverwort) and *Chlorella sp*. NC64A (green algae) [19, 20, 25, 40]. Specifically, a linear model was constructed between deep (*x*) and 1× (*y*) sequencing coverage, which was then used to extrapolate levels of DNA methylation and 95% confidence intervals (CI) from low sequence coverage species (Figure S10 and Tables S5).

### Genic DNA methylation analyses and metaplots

DNA methylation was estimated as weighted DNA methylation, which is the total number of aligned DNA methylated reads divide by the total number of methylated plus un-methylated reads. This metric of DNA methylation was estimated for each sequence context within coding regions. For *P. taeda* only high quality gene models were used, since low quality models cannot distinguish between pseudogenes and true protein coding genes. For genic metaplots, the gene body – start to stop codon – was divided into 20 windows. Additionally, for species with assembled and annotated genomes regions 1000 or 4000 bp upstream and downstream were divided into 20 windows. Weighted DNA methylation was calculated for each window. The mean weighted methylation for each window was then calculated for all genes and plotted in R v3.2.4 (https://www.r-project.org/).

### mCG-enrichment test

Sequence context enrichment for each gene was determined through a binomial test followed by Benjamini–Hochberg FDR [25, 26]. A context-specific background level of DNA methylation determined from the coding sequence was used as a threshold in determining significance. Genes were classified as mCG-enriched/gbM if they had reads mapping to at least 10 CG sites and a q-value <0.05 for mCG, and a q-value >0.05 for mCHG and mCHH.

## DECLARATIONS

## Acknowledgements

We thank Nathan Springer for comments and discussions. Kevin Tarner (UGA greenhouse), and Michael Wenzel and Ron Determann (Atlanta Botanical Gardens) for plant tissue. Brigitte T. Hofmeister for establishment and maintenance of the genome browser. We also thank the Georgia Genomics Facility (GGF) for sequencing. Computational resources were provided by the Georgia Advanced Computing Resource Center (GACRC). We thank Gane Ka-Shu Wong and the 1000 Plants initiative (1KP, onekp.com) for advanced access to transcript assemblies.

## Funding

The work conducted by the National Science Foundation (NSF) (MCB– 1402183) and by The Pew Charitable Trusts to R.J.S.

## Availability of data and materials

Genome browsers for all methylation data used in this paper are located at Plant Methylation DB (schmitzlab.genetics.uga.edu/plantmethylomes). Sequence data for MethylC-seq are located at the Gene Expression Omnibus, accession GSE81702.

## Authors’ contributions

Conceptualization: AJB, and RJS; Performed experiments: AJB, CEN, LJ, and NAR; Data Analysis: AJB, CEN, and LJ; Writing – Original Draft: AJB; Writing – Review and Editing: AJB, JL-M, and RJS; Resources: JL-M, and RJS. All authors read and approved the final manuscript.

## Competing interests

The authors declare that they have no competing interests.

## Ethics approval

Ethics approval was not needed for this study.

## SUPPLEMENTAL INFORMATION

**Figure S1. CMT proteins in green algae (*C. reinhardtii, Chlorella* NC64A, and *V. carteri*) might represent misidentified homologs. a**, A midpoint rooted gene tree constructed from a subset of species and green algae using protein sequences. Previously identified CMT homologs in *C. reinhardtii, Chlorella* NC64A, and *V. carteri* (JGI accession ids 190580, 52630, and 94056, respectively) have low amino acid sequence similarity to *A. thaliana* CMT compared to other green algae species (Table S1), which is reflected in long branches, especially for *C. reinhardtii* and *V. carteri*. Values on branches are raw branch lengths represented as amino acid substitutions per amino acid site. **b**, Protein structure of previously identified CMT homologs in *C. reinhardtii, Chlorella* NC64A, and *V. carteri* and those identified in green algae from the 1KP dataset. Reported CMTs in *C. reinhardtii* and *Chlorella* NC64A do not contain CHROMO domains, and the homolog in *V. carteri* does not contain any recognizable PFAM domains, however BAH, CHROMO and a DNA methylase domain can all be identified in green algae CMT homologs from the 1KP dataset.

**Figure S2. Phylogenetic relationships among CMTs in Viridiplantae**. CMTs are separated into four monophyletic clades based on bootstrap support and the relationship of *A. thaliana* CMTs: (i) the gbM-dependent CMT superclade with subclades CMT1, CMT3, ZMET and *A. trichopoda*; (ii) CMT2 and; (iii) homologous (hCMT) α and β. Values at nodes in represent bootstrap support from 1000 replicates, and the tree was rooted to the clade containing all green algae species.

**Figure S3. Syntenic relationships support a Whole Genome Duplication (WGD) event giving rise to CMT1 and CMT3 in eudicots and ZMET paralogs. a**, Synteny was determined using CoGe’s GEvo program, and is indicated by connected blocks. Synteny is more pronounced in some eudicots over others, which suggests sequence divergence following the shared WGD placed at the base of all eudicots [36]. **b**, Phylogenetic relationships of ZMETs in the Poaceae suggest WGD events are shared by several species and are species-specific as is the case for ZMET2 and ZMET5 in *Z. mays*. Colors following the tip labels indicate clades of paralogous ZMETs. **c**, Similarly to eudicots, WGD is supported by synteny upstream and downstream of ZMET paralogs.

**Figure S4. Presence and absence of CMTs and ZMETs in eudicots, and 736 monocots and monocots/commelinids, respectively. a**, Eudicot (basal, core, rosid, and asterid) species of plants possess different combinations of CMT1, CMT2, and CMT3. CMT3 was potentially loss from 46/262 (18%), and CMT1 is found in 106/262 (40%) of eudicot species sequenced by the 1KP Consortium. Species without CMT3 are predicted to have significantly reduced levels of gbM loci compared to eudicot species with CMT3. The presence of CMT1 in numerous species suggests a yet to be determined functional role of CMT1 in DNA methylation and/or chromatin modification. **b**, Similarly to eudicots, monocots and monocots/commelinids have different combinations of CMT2 and ZMET, which may reflect differences in genome structure, and DNA methylation and chromatin modification patterns.

**Figure S5. Metagene plots of DNA methylation across gene bodies**. DNA methylation levels within all full-length coding sequences or transcripts for additional species used in this study.

**Figure S6. MCG-enriched genes in species sister to angiosperms are rare 753 and not strongly conserved. a**, The proportion of mCG-enriched genes are variable across Embryophyta. However, lowest levels are seen in species null for CMT3 and that possess a non-orthologous CMT3 (white circles). Additionally, species that possess a CMT3 that has experienced elevated rates of evolution (ω) have a lower proportion of mCG-enriched genes (gray circles). **b**, The majority of mCG-enriched genes are orthologous to non-mCG-enriched genes in *A. thaliana* or have no hits to an *A. thaliana* gene based on an e-value of≤1E-06. However, *P. taeda* is an exception, which suggests some of the mCG-enriched genes are conserved to gbM genes in *A. thaliana*.

**Figure S7. MCG in genes of species sister to angiosperms are biased 764 towards extreme low or high levels**. Distributions of mCG across all genes with sufficient coverage for species with sequenced genomes (see Methods).

**Figure S8. Jumonji (jmjC) domain-containing gene family phylogeny**. The jmjC domain-containing family contains five monophyletic clades based on the location of *A. thaliana* genes. Only angiosperm sequences can be found within the clade containing *A. thaliana* IBM1. Scale bar represents nucleotide substitutions per site.

**Figure S9. SUVH4 and SUVH5/6 gene family phylogenies. a**, SUVH4 gene family approximately recapitulate species relationships, and angiosperm-specific monophyletic clades are not observed based on bootstrap support and the placement of *A. thaliana* SUVH4. **b**, However, Brassicaceae-specific monophyletic clades delineate SUVH5 and SUVH6, hence a homologous SUVH5/6 (hSUVH5/6) sequence is found in other Embryophyta. However, some nodes – especially those delineating monocot and monocot/commelinid hSUVH5/6 sequences – are weakly supported. **c**, Phylogenetic relationships support a Brassicacae-specific duplication event, which gave rise to SUVH5 and SUVH6. **d**, Reanalyzing monocot and monocot/commelinid hSUVH5/6 sequences increases bootstrap support delineating two monophyletic clades. This relationship is analogous to SUVH5 and SUVH6 in Brassicaceae, but encompasses all monocots and monocot/commelinids. Furthermore, Poaceae-specific monophyletic clades are observed within each of the monocot- and monocot/commelinid-specific monophyletic clades. Phylogenetic relationships support multiple duplication events in the monocots and monocot/commelinids. Values at nodes represent bootstrap support and scale bar represents nucleotide substitutions per site.

**Figure S10. A linear model to determine DNA methylation levels from low 793 sequence coverage WGBS**. A strong linear correlation is observed between DNA methylation levels at CG, CHG and CHH sites determined from low, subsampled and full WGBS coverage. A linear model was generated for each sequence context, which was used to extrapolate levels of DNA methylation from species with low WGBS coverage. Each data point represents a single plant species from [19, 20, 25, 40].

**Table S1. Taxonomic, sequence, and phylogenetic summary of sequences used in Fig. 1 and Supplementary Fig. 2**.

**Table S2. Best BLASTp hits of published green algae CMTs suggest mis-annotation compared to green algae CMTs identified in the current study**.

**Table S3. A summary of branch and branch-site tests implemented in PAML**.

**Table S4. Reduced (1×) and deep sequencing coverage estimates of DNA methylation levels from 34 Viridiplantae species**.

**Table S5. DNA methylation levels of species sequenced in this study and the levels predicted by a context-specific linear model**.

